# Differential Expression Analysis with InMoose, the Integrated Multi-Omic Open-Source Environment in Python

**DOI:** 10.1101/2024.11.14.623578

**Authors:** Maximilien Colange, Guillaume Appé, Léa Meunier, Solène Weill, Akpéli Nordor, Abdelkader Behdenna

## Abstract

We present the differential expression features of InMoose, a Python implementation of R tools *limma, edgeR*, and *DESeq2*. We experimentally show that InMoose stands as a drop-in replacement for those tools, with nearly identical results. This ensures reproducibility when interfacing both languages in bioinformatic pipelines. InMoose is an open source software released under the GPL3 license, available at www.github.com/epigenelabs/inmoose and https://inmoose.readthedocs.io.

## Introduction

Differential expression analysis (DEA) is a prominent technique for the analysis of high-dimensional biomolecular data that helps scientists discover genetic features associated with studied phenotypes. Despite the historical prevalence of R, Python is the standard language for techniques that are becoming essential parts of bioinformatic pipelines: *e*.*g*. data science, machine learning, or artificial intelligence. More generally, Python’s wide adoption across a variety of domains increases its versatility and eases tool integration. While recent tools are developed directly in Python (*omicverse* ^1^, *scverse* ^2^…), state-of-the-art DEA tools are available in R only. To our knowledge, *pydeseq2* ^3^ is currently the sole Python tool to offer generalized linear model-based DEA.

Continuing our previous effort ^4^, we have ported into InMoose three state-of-the-art DEA tools: *limma* ^5^, *edgeR* ^6^ and *DESeq2* ^7^. We focus on providing drop-in replacements for the R tools, allowing practitioners to easily navigate between R and Python ecosystems while retaining reproducibility and comparability across languages.

## Implementation

InMoose implements bulk transcriptomic DEA methods from three state-of-the-art tools:

- *limma* ^5^ is an R package using empirical Bayesian methods and linear models for assessing differential gene expression. Initially targeting microarray technologies, it applies to other technologies yielding data with similar statistical behavior.
- *edgeR* ^6^ is an R package also based on empirical Bayesian methods and linear models for the analysis of gene expression data. Its models are specifically geared towards next-generation RNA-sequencing (RNA-Seq) data.
- *DESeq2* ^7^ is another R package based on empirical Bayesian methods and linear models for the analysis of gene expression data, also geared towards RNA-Seq data. In addition to DEA, *DESeq2* features are widely used for data normalization.

Rather than re-implementing the methods from scratch, we ported the R code to Python, following two goals:

- provide a drop-in replacement for R tools, with similar – if not identical – results to ensure reproducibility and avoid regressions during tool migration;
- capitalize on existing implementations: implementation details do matter, yet they are seldom described in publications or documentation. When porting code, our primary source of knowledge is the tool source code.

To illustrate this, we compare the results obtained with InMoose to those obtained with *limma, edgeR* and *DESeq2*, and with *pydeseq2* ^3^, an alternative Python implementation of *DESeq2*.

### Experimental Data and Setup

We selected 12 microarray and 7 RNA-Seq datasets from GEO ^8^, each featuring both healthy and tumor tissue samples:

- microarray
  - colorectal cancer: GSE20916, GSE23194, GSE37364, GSE4183, GSE44076, GSE62932
  - ovary cancer: GSE18520, GSE23391, GSE36668, GSE38666, GSE52037, GSE54388
- RNA-Seq
  - brain cancer: GSE147352, GSE148389, GSE205512, GSE205590
  - breast cancer: GSE174339,
  - ovary cancer: GSE212991, GSE254461

The experiments were run with Python 3.11.4 and R 4.2.2, with the following tool versions:

- InMoose 0.7.3
- *limma* 3.54.2
- *edgeR* 3.40.2
- *DESeq2* 1.38.3
- *pydeseq2* 0.4.11

### Comparison with *limma*

On each microarray dataset, we computed the log-fold-changes (LFC) between the cancer and healthy samples groups, with the InMoose limma and the *limma* pipelines. Figure 1A compares the LFC from both tools, showing the distribution of their absolute differences and Pearson correlation on each dataset.

**Figure 1.**
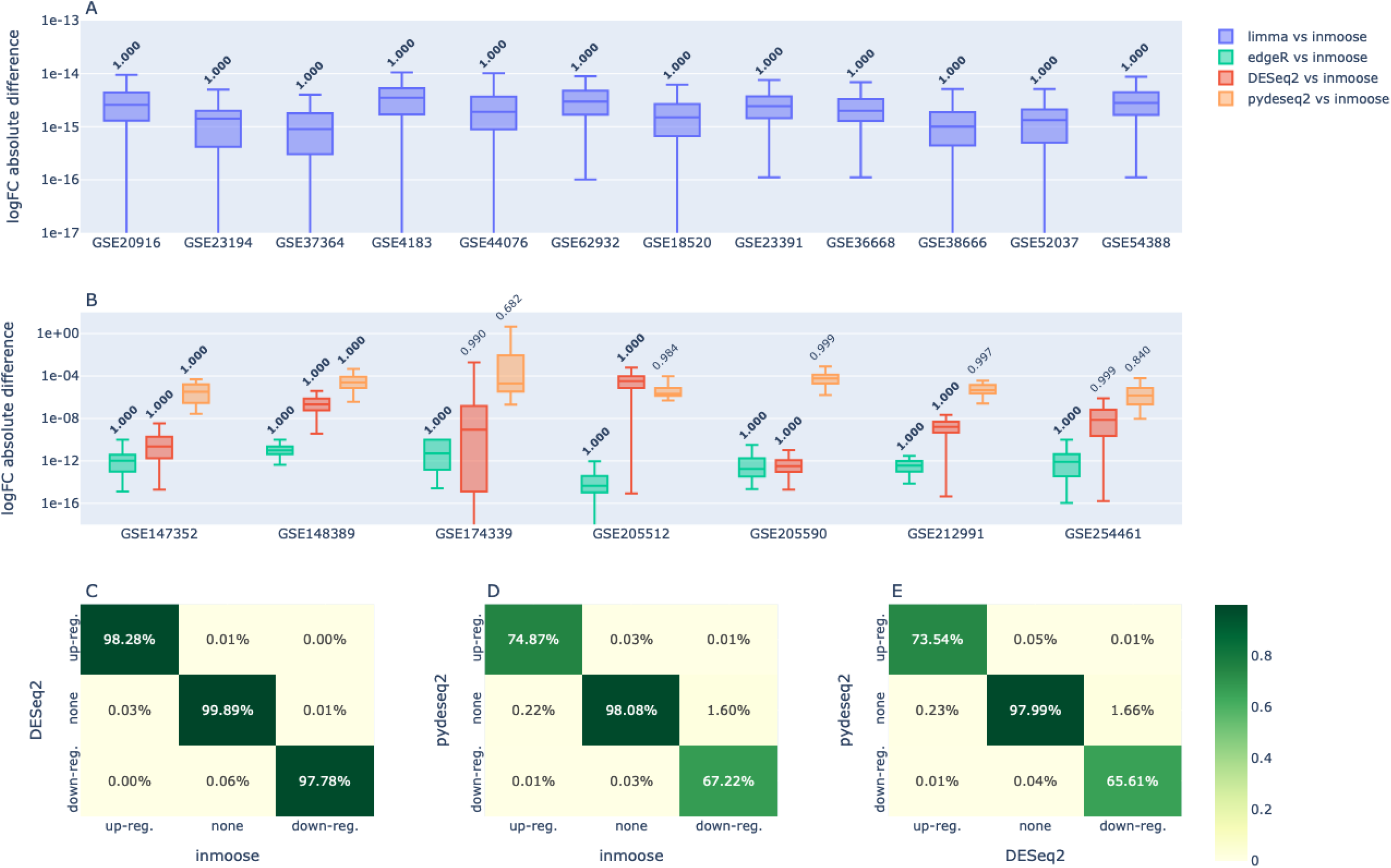
Comparison of InMoose with *limma, edgeR, DESeq2* and *pydeseq2*. **A**. Absolute differences of log-fold-change between *limma* and InMoose, and Pearson correlations. Boxes represent the median, and first and third quartiles, whiskers represent the 2.5% and 97.5% quantiles. **B**. Absolute differences of log-fold-change between *edgeR, DESeq2, pydeseq2* and InMoose, and Pearson correlations. Boxes represent the median, and first and third quartiles, whiskers represent the 2.5% and 97.5% quantiles. **C**. Significantly differentially expressed genes (padj < 0.05, |LFC| > 2) according to InMoose and *DESeq2*. Percentages averaged across the 7 datasets. **D**. Significantly differentially expressed genes (padj < 0.05, |LFC| > 2) according to InMoose and *pydeseq2*. Percentages averaged across the 7 datasets. **E**. Significantly differentially expressed genes (padj < 0.05, |LFC| > 2) according to *DESeq2* and *pydeseq2*. Percentages averaged across the 7 datasets.

The results obtained by both tools are almost identical, with a Pearson correlation of 100% on all datasets, and very low LFC absolute differences – ranging below 1e-14. This shows that InMoose is a solid drop-in replacement for *limma*, with very low risks of regression or divergence in the results obtained.

### Comparison with *edgeR*

On each RNA-Seq dataset, we computed the log-fold-change (LFC) between the cancer and healthy samples groups, with the InMoose edgepy LRT and the *edgeR* LRT pipelines. Figure 1B compares the LFC obtained from both tools, showing the distribution of their absolute differences and Pearson correlation on each dataset.

The results obtained by both tools are almost identical, with a Pearson correlation of 100% on all datasets, and very low LFC absolute differences – ranging below 1e-8. This shows that InMoose is a solid drop-in replacement for *edgeR*, with very low risks of regression or divergence in the results obtained.

### Comparison with *DESeq2* and *pydeseq*

On each RNA-Seq dataset, we computed the log-fold-change (LFC) between the cancer and healthy samples groups, with the InMoose deseq2, the *DESeq2* and the *pydeseq2* pipelines. Figure 1B compares the LFC obtained from InMoose deseq2 and *DESeq2*, on the one hand, and from InMoose deseq2 and *pydeseq2*, on the other hand. The figure shows the distribution of the LFC absolute differences and Pearson correlation on each dataset between both pairs of tools.

The results obtained by InMoose deseq2 and *DESeq2* are very similar, with Pearson correlations above 99%, and most LFC absolute differences below 1e-4 – although larger differences occur on datasets where LFC are least correlated. Figure 1C confirmed this high similarity, since InMoose and *DESeq2* agree on differentially expressed genes.

In contrast, the results obtained by *pydeseq2* are more heterogeneous. Although the LFC correlations with InMoose are overall high (above 98% for 5 datasets out of 7, above 60% for the remaining 2), the LFC absolute differences between InMoose and *pydeseq2* are significantly higher and with higher extremal values than between InMoose and *DESeq2*. While the median deviation remains below 1e-4 on all datasets, such fluctuations may significantly impact threshold- or order-based analysis methods. Indeed, the differentially expressed genes identified by *pydeseq2* differ significantly from those found by InMoose and *DESeq2*, as shown on Figure 1D and 1E, respectively.

These results show that InMoose and *DESeq2* have very similar results, both quantitatively and qualitatively, while *pydeseq2* qualitative results diverge. We conclude that InMoose should be preferred when migrating *DESeq2* pipelines from R to Python, to minimize the impact on results and to preserve comparability of R-based and Python-based pipelines.

## Conclusion

In conclusion, InMoose is a reliable package for microarray and RNA-Seq DEA. The similarity of its results with those of *limma, edgeR* and *DESeq2* makes it the solution of choice to ensure comparability of Python-based and R-based analyses. InMoose addresses a gap in the current Python ecosystem while minimizing the reproducibility risks associated with the migration from R to Python.

## References

1. Zeng, Z. et al. OmicVerse: a framework for bridging and deepening insights across bulk and single-cell sequencing. Nat. Commun. 15, 5983 (2024).

2. Virshup, I. et al. The scverse project provides a computational ecosystem for single-cell omics data analysis. Nat. Biotechnol. 41, 604–606 (2023).

3. Muzellec, B., Teleńczuk, M., Cabeli, V. & Andreux, M. PyDESeq2: a python package for bulk RNA-seq differential expression analysis. Bioinformatics 39, btad547 (2023).

4. Behdenna, A. et al. pyComBat, a Python tool for batch effects correction in high-throughput molecular data using empirical Bayes methods. BMC Bioinformatics 24, 459 (2023).

5. Ritchie, M. E. et al. limma powers differential expression analyses for RNA-sequencing and microarray studies. Nucleic Acids Res. 43, e47–e47 (2015).

6. Chen, Y., Lun, A. T. L. & Smyth, G. K. From reads to genes to pathways: differential expression analysis of RNA-Seq experiments using Rsubread and the edgeR quasi-likelihood pipeline. Preprint at 10.12688/f1000research.8987.2 (2016).

7. Love, M. I., Huber, W. & Anders, S. Moderated estimation of fold change and dispersion for RNA-seq data with DESeq2. Genome Biol. 15, 550 (2014).

8. Barrett, T. et al. NCBI GEO: archive for functional genomics data sets—update. Nucleic Acids Res. 41, D991–D995 (2012).

