## Supplementary material for "Differential Expression Analysis with InMoose, the Integrated Multi-Omic Open-Source Environment in Python": README.md

# InMoose Comparison with limma, edgeR, DESeq2 and pydeseq2

This supplementary material contains the experimental data and scripts to
reproduce the experiments reported in the paper "Differential Expression
Analysis with InMoose, the Integrated Multi-Omic Open-Source Environment in
Python", including re-generating the figure of the paper.

## Content

`installR.R`, `requirements.txt`, `run.py`, `run_limma.R`, `run_edgeR.R`,
`run_DESeq2.R` are script files to reproduce the experiments of the paper.

`GSE*` folders contain the clinical and molecular data of the corresponding
dataset.

`*.csv` files contain the experimental results we obtain and report in the
paper.

## Reproducing the Experiments

Make sure Python and R are installed on the machine.

Install Python dependencies: `pip install -r requirements.txt`.
Install R dependencies: `Rscript installR.R`.

Run the script: `python run.py`.
This scripts will call all Python and R tools for you, and regenerates the
figure of the paper.
