## Supplementary material for "Differential Expression Analysis with InMoose, the Integrated Multi-Omic Open-Source Environment in Python": run.py

```
import subprocess
import time

import numpy as np
import pandas as pd
import patsy
from inmoose.deseq2 import DESeq, DESeqDataSet
from inmoose.edgepy import DGEList, glmLRT
from inmoose.limma import contrasts_fit, eBayes, lmFit, makeContrasts, topTable
from pydeseq2.dds import DeseqDataSet
from pydeseq2.ds import DeseqStats
from scipy.stats import pearsonr

datasets = {
    "GSE147352": {
        "factors": ["metapipeline_sample_type"],
        "contrast": ["metapipeline_sample_type", "Normal tissue", "Primary tumor"],
    },
    "GSE148389": {
        "factors": ["metapipeline_sample_type"],
        "contrast": ["metapipeline_sample_type", "Normal tissue", "Primary tumor"],
    },
    "GSE174339": {
        "factors": ["metapipeline_sample_type"],
        "contrast": ["metapipeline_sample_type", "Normal tissue", "Primary tumor"],
    },
    "GSE205512": {
        "factors": ["metapipeline_sample_type"],
        "contrast": ["metapipeline_sample_type", "Normal tissue", "Primary tumor"],
    },
    "GSE205590": {
        "factors": ["metapipeline_sample_type"],
        "contrast": ["metapipeline_sample_type", "Normal tissue", "Primary tumor"],
    },
    "GSE212991": {
        "factors": ["metapipeline_sample_type"],
        "contrast": ["metapipeline_sample_type", "Normal tissue", "Primary tumor"],
    },
    "GSE254461": {
        "factors": ["metapipeline_sample_type"],
        "contrast": ["metapipeline_sample_type", "Normal tissue", "Primary tumor"],
    },
}

datasets_microarray = {
    "GSE20916": {
        "factors": ["metapipeline_sample_type"],
        "contrast": ["metapipeline_sample_type", "Normal tissue", "Primary tumor"],
    },
    "GSE23194": {
        "factors": ["metapipeline_sample_type"],
        "contrast": ["metapipeline_sample_type", "Normal tissue", "Primary tumor"],
    },
    "GSE37364": {
        "factors": ["metapipeline_sample_type"],
        "contrast": ["metapipeline_sample_type", "Normal tissue", "Primary tumor"],
    },
    "GSE4183": {
        "factors": ["metapipeline_sample_type"],
        "contrast": ["metapipeline_sample_type", "Normal tissue", "Primary tumor"],
    },
    "GSE44076": {
        "factors": ["metapipeline_sample_type"],
        "contrast": ["metapipeline_sample_type", "Normal tissue", "Primary tumor"],
    },
    "GSE62932": {
        "factors": ["metapipeline_sample_type"],
        "contrast": ["metapipeline_sample_type", "Normal tissue", "Primary tumor"],
    },
    "GSE18520": {
        "factors": ["metapipeline_sample_type"],
        "contrast": ["metapipeline_sample_type", "Normal tissue", "Primary tumor"],
    },
    "GSE23391": {
        "factors": ["metapipeline_sample_type"],
        "contrast": ["metapipeline_sample_type", "Normal tissue", "Primary tumor"],
    },
    "GSE36668": {
        "factors": ["metapipeline_sample_type"],
        "contrast": ["metapipeline_sample_type", "Normal tissue", "Primary tumor"],
    },
    "GSE38666": {
        "factors": ["metapipeline_sample_type"],
        "contrast": ["metapipeline_sample_type", "Normal tissue", "Primary tumor"],
    },
    "GSE52037": {
        "factors": ["metapipeline_sample_type"],
        "contrast": ["metapipeline_sample_type", "Normal tissue", "Primary tumor"],
    },
    "GSE54388": {
        "factors": ["metapipeline_sample_type"],
        "contrast": ["metapipeline_sample_type", "Normal tissue", "Primary tumor"],
    },
}


def run_deseq():
    times = {}
    for ds, config in datasets.items():
        factors = config["factors"]
        contrast = config["contrast"]
        print(f"analyzing {ds}")
        molecular = pd.read_csv(f"{ds}/gene_expression_df.csv", index_col=0)
        clinical = pd.read_csv(f"{ds}/clinical_df.csv", index_col=0)
        clinical = clinical.loc[molecular.columns, :]

        # inmoose.deseq2
        print("running inmoose.deseq")
        start = time.time()
        dds = DESeqDataSet(molecular.T, clinical, design=f"~ 1 + {'+'.join(factors)}")
        dds = DESeq(dds)
        inmoose_res = dds.results(contrast=contrast)
        end = time.time()
        times[ds] = {"inmoose": end - start}
        inmoose_res.to_csv(f"{ds}_inmoose.csv")

        # pydeseq2
        print("running pydeseq2")
        start = time.time()
        data = DeseqDataSet(
            counts=molecular.T, metadata=clinical, design_factors=factors
        )
        data.deseq2()
        contrast2 = [c.replace("_", "-") for c in contrast]
        stats = DeseqStats(data, contrast=contrast2, quiet=True)
        stats.summary()
        pydeseq2_res = stats.results_df
        end = time.time()
        times[ds]["pydeseq2"] = end - start
        pydeseq2_res.to_csv(f"{ds}_pydeseq2.csv")

        # DESeq2
        # call a R script, that loads the csv, runs R DESeq2 and saves results to file
        start = time.time()
        with subprocess.Popen(
            ["Rscript", "run_DESeq2.R", ds, ",".join(factors), ",".join(contrast)]
        ):
            pass
        end = time.time()
        times[ds]["R"] = end - start

    print(times)


def run_edger():
    times = {}
    for ds, config in datasets.items():
        factors = config["factors"]
        contrast = config["contrast"]
        print(f"analyzing {ds}")
        molecular = pd.read_csv(f"{ds}/gene_expression_df.csv", index_col=0)
        clinical = pd.read_csv(f"{ds}/clinical_df.csv", index_col=0)
        clinical = clinical.loc[molecular.columns, :]

        # inmoose.edgepy
        print("running inmoose.edgeR")
        start = time.time()
        d = DGEList(counts=molecular)
        d.design = patsy.dmatrix(f"~1 + {'+'.join(factors)}", data=clinical)
        print("estimating common disp")
        d = d.estimateGLMCommonDisp()
        # LRT
        print("computing LRT")
        edgepy_lrt = glmLRT(d.glmFit())
        # QL FTest
        print("computing QL")
        # edgepy_ql = glmQLFTest(d.glmQLFit())
        # exactTest
        print("computing exact")
        # edgepy_exact = exactTest(d)
        end = time.time()
        times[ds] = {"inmoose": end - start}
        edgepy_lrt.to_csv(f"{ds}_edgepy_lrt.csv")
        # edgepy_ql.to_csv(f"{ds}_edgepy_ql.csv")
        # edgepy_exact.to_csv(f"{ds}_edgepy_exact.csv")

        # edgeR
        # call a R script, that loads the csv, runs R edgeR and saves results to file
        start = time.time()
        with subprocess.Popen(
            ["Rscript", "run_edgeR.R", ds, ",".join(factors), ",".join(contrast)]
        ):
            pass
        end = time.time()
        times[ds]["R"] = end - start

    print(times)


def run_limma():
    times = {}
    for ds, config in datasets_microarray.items():
        factors = config["factors"]
        contrast = config["contrast"]
        print(f"analyzing {ds}")
        molecular = pd.read_csv(f"{ds}/gene_expression_df.csv", index_col=0)
        clinical = pd.read_csv(f"{ds}/clinical_df.csv", index_col=0)
        clinical = clinical.loc[molecular.columns, :]

        nc1 = contrast[1].replace(" ", "_")
        nc2 = contrast[2].replace(" ", "_")
        clinical[contrast[0]] = (
            clinical[contrast[0]].replace(contrast[1], nc1).replace(contrast[2], nc2)
        )
        contrast[1] = nc1
        contrast[2] = nc2

        # inmoose.limma
        print("running inmoose.limma")
        start = time.time()
        design = patsy.dmatrix(f"~0 + {'+'.join(factors)}", data=clinical)
        fit1 = lmFit(molecular, design)
        tt = f"{contrast[0]}[{contrast[2]}] - {contrast[0]}[{contrast[1]}]"
        contrast_matrix = makeContrasts(tt, levels=design)
        data_fit_con = contrasts_fit(fit1, contrast_matrix)
        fit1_ebay = eBayes(data_fit_con)
        top_table = topTable(fit1_ebay, number=np.inf, coef=tt)
        end = time.time()
        times[ds] = {"inmoose": end - start}
        top_table.to_csv(f"{ds}_pylimma.csv")

        # limma
        # call a R script, that loads the csv, runs R edgeR and saves results to file
        start = time.time()
        with subprocess.Popen(
            ["Rscript", "run_limma.R", ds, ",".join(factors), ",".join(contrast)]
        ):
            pass
        end = time.time()
        times[ds]["R"] = end - start

    print(times)


def plot_barplots():
    import plotly.graph_objects as go
    from plotly.subplots import make_subplots

    quantiles = [0.025, 0.25, 0.5, 0.75, 0.975]

    fig = make_subplots(
        rows=3,
        cols=3,
        specs=[
            [{"type": "box", "colspan": 3}, None, None],
            [{"type": "box", "colspan": 3}, None, None],
            [{}, {}, {}],
        ],
        subplot_titles=["A", "B", "C", "D", "E"],
        vertical_spacing=0.1,
        horizontal_spacing=0.1,
    )
    # align subplot titles to the left
    fig.update_annotations(x=0.01)
    fig.update_annotations(selector=3, xref="x4 domain", x=0.03)
    fig.update_annotations(selector=4, xref="x5 domain", x=0.03)

    # limma
    data = pd.DataFrame(columns=["median", "q1", "q3", "lf", "uf", "pearson"])
    for ds in datasets_microarray.keys():
        pylimma = pd.read_csv(f"{ds}_pylimma.csv", index_col=0)
        limma = pd.read_csv(f"{ds}_limma.csv", index_col=0)

        diff = pylimma["log2FoldChange"] - limma["logFC"]
        denom = np.maximum(pylimma["log2FoldChange"].abs(), limma["logFC"].abs())
        assert all(diff[denom == 0] == 0)
        denom[denom == 0] = 1e-10
        diff = diff.abs()  # / denom

        data.loc[ds] = (
            np.quantile(diff, quantiles[2]),
            np.quantile(diff, quantiles[1]),
            np.quantile(diff, quantiles[3]),
            np.quantile(diff, quantiles[0]),
            np.quantile(diff, quantiles[4]),
            pearsonr(pylimma["log2FoldChange"], limma["logFC"]).statistic,
        )

    fig.add_trace(
        go.Box(
            x=data.index,
            median=data["median"],
            q1=data["q1"],
            q3=data["q3"],
            lowerfence=data["lf"],
            upperfence=data["uf"],
            boxpoints=False,
            name="limma vs inmoose",
            offsetgroup="limma vs inmoose",
        ),
        1,
        1,
    )
    for ds in data.index:
        t = f"{data.loc[ds, 'pearson']:.3f}"
        fig.add_annotation(
            x=ds,
            y=np.log10(data.loc[ds, "uf"]),
            text=t,
            row=1,
            col=1,
            showarrow=False,
            xshift=-1,
            yshift=18,
            font={"size": 10, "weight": "bold" if t == "1.000" else "normal"},
            textangle=315,
        )

    # edgeR
    data = pd.DataFrame(columns=["median", "q1", "q3", "lf", "uf", "pearson"])
    for ds in datasets.keys():
        edgepy = pd.read_csv(f"{ds}_edgepy_lrt.csv", index_col=0)
        edger = pd.read_csv(f"{ds}_edgeR.csv", index_col=0)
        assert all(edgepy.index == edger.index)
        # edgepy_nan = edgepy["log2FoldChange"].isna()
        # edger_nan = edger["logFC"].isna()
        # edgepy = edgepy.loc[~(edgepy_nan | edger_nan)]
        # edger = edger.loc[~(edgepy_nan | edger_nan)]
        edgepy.loc[edgepy["log2FoldChange"] == 0, "logFoldChange"] = 1e-10
        edger.loc[edger["logFC"] == 0, "logFC"] = 1e-10

        diff = edgepy["log2FoldChange"] - edger["logFC"]
        denom = np.maximum(edgepy["log2FoldChange"].abs(), edger["logFC"].abs())
        assert all(diff[denom == 0] == 0)
        denom[denom == 0] = 1e-10
        diff = diff.abs()  # / denom

        data.loc[ds] = (
            np.quantile(diff, quantiles[2]),
            np.quantile(diff, quantiles[1]),
            np.quantile(diff, quantiles[3]),
            np.quantile(diff, quantiles[0]),
            np.quantile(diff, quantiles[4]),
            pearsonr(edgepy["log2FoldChange"], edger["logFC"]).statistic,
        )

    fig.add_trace(
        go.Box(
            x=data.index,
            median=data["median"],
            q1=data["q1"],
            q3=data["q3"],
            lowerfence=data["lf"],
            upperfence=data["uf"],
            boxpoints=False,
            name="edgeR vs inmoose",
            offsetgroup="edgeR vs inmoose",
        ),
        2,
        1,
    )
    for ds in data.index:
        t = f"{data.loc[ds, 'pearson']:.3f}"
        fig.add_annotation(
            x=ds,
            y=np.log10(data.loc[ds, "uf"]),
            text=t,
            row=2,
            col=1,
            showarrow=False,
            xshift=-33,
            yshift=18,
            font={"size": 10, "weight": "bold" if t == "1.000" else "normal"},
            textangle=315,
        )

    # DESeq2
    data = pd.DataFrame(columns=["median", "q1", "q3", "lf", "uf", "pearson"])
    hm = {}
    for ds in datasets.keys():
        inmoose = pd.read_csv(f"{ds}_inmoose.csv", index_col=0)
        deseq = pd.read_csv(f"{ds}_deseq2.csv", index_col=0)
        inmoose_nan = inmoose["log2FoldChange"].isna()
        deseq_nan = deseq["log2FoldChange"].isna()
        inmoose = inmoose.loc[~(inmoose_nan | deseq_nan)]
        deseq = deseq.loc[~(inmoose_nan | deseq_nan)]
        assert all(inmoose.index == deseq.index)

        diff = inmoose["log2FoldChange"] - deseq["log2FoldChange"]
        denom = np.maximum(
            inmoose["log2FoldChange"].abs(), deseq["log2FoldChange"].abs()
        )
        assert all(diff[denom == 0] == 0)
        denom[denom == 0] = 1e-10
        diff = diff.abs()  # / denom

        data.loc[ds] = (
            np.quantile(diff, quantiles[2]),
            np.quantile(diff, quantiles[1]),
            np.quantile(diff, quantiles[3]),
            np.quantile(diff, quantiles[0]),
            np.quantile(diff, quantiles[4]),
            pearsonr(inmoose["log2FoldChange"], deseq["log2FoldChange"]).statistic,
        )

        mat = pd.DataFrame(
            columns=["inmoose up", "inmoose neutral", "inmoose down"],
            index=["deseq up", "deseq neutral", "deseq down"],
        )
        inmoose_up = set(
            inmoose.index[(inmoose["log2FoldChange"] > 2) & (inmoose["padj"] < 0.05)]
        )
        inmoose_down = set(
            inmoose.index[(inmoose["log2FoldChange"] < -2) & (inmoose["padj"] < 0.05)]
        )
        inmoose_neutral = set(inmoose.index) - inmoose_up - inmoose_down

        deseq_up = set(
            deseq.index[(deseq["log2FoldChange"] > 2) & (deseq["padj"] < 0.05)]
        )
        # print(ds, len(deseq_up), len(inmoose_up))
        deseq_down = set(
            deseq.index[(deseq["log2FoldChange"] < -2) & (deseq["padj"] < 0.05)]
        )
        deseq_neutral = set(deseq.index) - deseq_up - deseq_down

        a = [inmoose_up, inmoose_neutral, inmoose_down]
        b = [deseq_up, deseq_neutral, deseq_down]
        for i in range(3):
            for j in range(3):
                mat.iloc[i, j] = len(b[i] & a[j]) / len(b[i] | a[j])
        hm[ds] = mat

    fig.add_trace(
        go.Box(
            x=data.index,
            median=data["median"],
            q1=data["q1"],
            q3=data["q3"],
            lowerfence=data["lf"],
            upperfence=data["uf"],
            boxpoints=False,
            name="DESeq2 vs inmoose",
            offsetgroup="DESeq2 vs inmoose",
        ),
        2,
        1,
    )
    for ds in data.index:
        t = f"{data.loc[ds, 'pearson']:.3f}"
        fig.add_annotation(
            x=ds,
            y=np.log10(data.loc[ds, "uf"]),
            text=t,
            row=2,
            col=1,
            showarrow=False,
            xshift=-1,
            yshift=18,
            font={"size": 10, "weight": "bold" if t == "1.000" else "normal"},
            textangle=315,
        )

    mean_hm = pd.DataFrame(
        columns=["inmoose up", "inmoose neutral", "inmoose down"],
        index=["deseq down", "deseq neutral", "deseq up"],
    )
    for c in mean_hm.columns:
        for i in mean_hm.index:
            mean_hm.loc[i, c] = np.mean([h.loc[i, c] for h in hm.values()])
    fig.add_trace(
        go.Heatmap(
            z=mean_hm,
            text=[
                [f"{100*mean_hm.iloc[i,j]:.2f}%" for j in range(mean_hm.shape[1])]
                for i in range(mean_hm.shape[0])
            ],
            texttemplate="%{text}",
            x=["up-reg.", "none", "down-reg."],
            y=["down-reg.", "none", "up-reg."],
            coloraxis="coloraxis",
        ),
        3,
        1,
    )

    # Owkin
    data = pd.DataFrame(columns=["median", "q1", "q3", "lf", "uf", "pearson"])
    hm = {}
    for ds in datasets.keys():
        inmoose = pd.read_csv(f"{ds}_inmoose.csv", index_col=0)
        pydeseq = pd.read_csv(f"{ds}_pydeseq2.csv", index_col=0)
        inmoose_nan = inmoose["log2FoldChange"].isna()
        pydeseq_nan = pydeseq["log2FoldChange"].isna()
        inmoose = inmoose.loc[~(inmoose_nan | pydeseq_nan)]
        pydeseq = pydeseq.loc[~(inmoose_nan | pydeseq_nan)]
        assert all(inmoose.index == pydeseq.index)

        diff = inmoose["log2FoldChange"] - pydeseq["log2FoldChange"]
        denom = np.maximum(
            inmoose["log2FoldChange"].abs(), pydeseq["log2FoldChange"].abs()
        )
        assert all(diff[denom == 0] == 0)
        denom[denom == 0] = 1e-10
        diff = diff.abs()  # / denom

        data.loc[ds] = (
            np.quantile(diff, quantiles[2]),
            np.quantile(diff, quantiles[1]),
            np.quantile(diff, quantiles[3]),
            np.quantile(diff, quantiles[0]),
            np.quantile(diff, quantiles[4]),
            pearsonr(inmoose["log2FoldChange"], pydeseq["log2FoldChange"]).statistic,
        )

        mat = pd.DataFrame(
            columns=["inmoose up", "inmoose neutral", "inmoose down"],
            index=["pydeseq up", "pydeseq neutral", "pydeseq down"],
        )
        inmoose_up = set(
            inmoose.index[(inmoose["log2FoldChange"] > 2) & (inmoose["padj"] < 0.05)]
        )
        inmoose_down = set(
            inmoose.index[(inmoose["log2FoldChange"] < -2) & (inmoose["padj"] < 0.05)]
        )
        inmoose_neutral = set(inmoose.index) - inmoose_up - inmoose_down

        pydeseq_up = set(
            pydeseq.index[(pydeseq["log2FoldChange"] > 2) & (pydeseq["padj"] < 0.05)]
        )
        pydeseq_down = set(
            pydeseq.index[(pydeseq["log2FoldChange"] < -2) & (pydeseq["padj"] < 0.05)]
        )
        pydeseq_neutral = set(pydeseq.index) - pydeseq_up - pydeseq_down
        # print(ds, len(pydeseq_up), len(inmoose_up))

        a = [inmoose_up, inmoose_neutral, inmoose_down]
        b = [pydeseq_up, pydeseq_neutral, pydeseq_down]
        for i in range(3):
            for j in range(3):
                mat.iloc[i, j] = len(b[i] & a[j]) / len(b[i] | a[j])
        hm[ds] = mat

    fig.add_trace(
        go.Box(
            x=data.index,
            median=data["median"],
            q1=data["q1"],
            q3=data["q3"],
            lowerfence=data["lf"],
            upperfence=data["uf"],
            boxpoints=False,
            name="pydeseq2 vs inmoose",
            offsetgroup="pydeseq2 vs inmoose",
        ),
        2,
        1,
    )
    for ds in data.index:
        t = f"{data.loc[ds, 'pearson']:.3f}"
        fig.add_annotation(
            x=ds,
            y=np.log10(data.loc[ds, "uf"]),
            text=t,
            row=2,
            col=1,
            showarrow=False,
            xshift=31,
            yshift=18,
            font={"size": 10, "weight": "bold" if t == "1.000" else "normal"},
            textangle=315,
        )
    mean_hm = pd.DataFrame(
        columns=["inmoose up", "inmoose neutral", "inmoose down"],
        index=["pydeseq down", "pydeseq neutral", "pydeseq up"],
    )
    for c in mean_hm.columns:
        for i in mean_hm.index:
            mean_hm.loc[i, c] = np.mean([h.loc[i, c] for h in hm.values()])
    fig.add_trace(
        go.Heatmap(
            z=mean_hm,
            text=[
                [f"{100*mean_hm.iloc[i,j]:.2f}%" for j in range(mean_hm.shape[1])]
                for i in range(mean_hm.shape[0])
            ],
            texttemplate="%{text}",
            x=["up-reg.", "none", "down-reg."],
            y=["down-reg.", "none", "up-reg."],
            coloraxis="coloraxis",
        ),
        3,
        2,
    )

    # Owkin vs. DESeq (heatmap only)
    hm = {}
    for ds in datasets.keys():
        deseq = pd.read_csv(f"{ds}_deseq2.csv", index_col=0)
        pydeseq = pd.read_csv(f"{ds}_pydeseq2.csv", index_col=0)
        deseq_nan = deseq["log2FoldChange"].isna()
        pydeseq_nan = pydeseq["log2FoldChange"].isna()
        deseq = deseq.loc[~(deseq_nan | pydeseq_nan)]
        pydeseq = pydeseq.loc[~(deseq_nan | pydeseq_nan)]
        assert all(deseq.index == pydeseq.index)

        mat = pd.DataFrame(
            columns=["deseq up", "deseq neutral", "deseq down"],
            index=["pydeseq up", "pydeseq neutral", "pydeseq down"],
        )
        deseq_up = set(
            deseq.index[(deseq["log2FoldChange"] > 2) & (deseq["padj"] < 0.05)]
        )
        deseq_down = set(
            deseq.index[(deseq["log2FoldChange"] < -2) & (deseq["padj"] < 0.05)]
        )
        deseq_neutral = set(deseq.index) - deseq_up - deseq_down

        pydeseq_up = set(
            pydeseq.index[(pydeseq["log2FoldChange"] > 2) & (pydeseq["padj"] < 0.05)]
        )
        pydeseq_down = set(
            pydeseq.index[(pydeseq["log2FoldChange"] < -2) & (pydeseq["padj"] < 0.05)]
        )
        pydeseq_neutral = set(pydeseq.index) - pydeseq_up - pydeseq_down

        a = [deseq_up, deseq_neutral, deseq_down]
        b = [pydeseq_up, pydeseq_neutral, pydeseq_down]
        for i in range(3):
            for j in range(3):
                mat.iloc[i, j] = len(b[i] & a[j]) / len(b[i] | a[j])
        hm[ds] = mat

    mean_hm = pd.DataFrame(
        columns=["deseq up", "deseq neutral", "deseq down"],
        index=["pydeseq down", "pydeseq neutral", "pydeseq up"],
    )
    for c in mean_hm.columns:
        for i in mean_hm.index:
            mean_hm.loc[i, c] = np.mean([h.loc[i, c] for h in hm.values()])
    fig.add_trace(
        go.Heatmap(
            z=mean_hm,
            text=[
                [f"{100*mean_hm.iloc[i,j]:.2f}%" for j in range(mean_hm.shape[1])]
                for i in range(mean_hm.shape[0])
            ],
            texttemplate="%{text}",
            x=["up-reg.", "none", "down-reg."],
            y=["down-reg.", "none", "up-reg."],
            coloraxis="coloraxis",
        ),
        3,
        3,
    )

    fig.update_layout(
        boxmode="group",
        autosize=False,
        height=910,
        width=1300,
        coloraxis={
            "colorscale": "ylgn",
            "colorbar": {"len": 0.35, "yanchor": "bottom", "y": -0.07},
        },
    )
    fig.update_yaxes(
        range=[-17, -13],
        type="log",
        row=1,
        tickmode="array",
        tickvals=[10**-i for i in range(13, 18)],
        ticktext=[f"{10**-i:.0e}" for i in range(13, 18)],
        title="logFC absolute difference",
    )
    fig.update_yaxes(
        range=[-18, 2],
        type="log",
        row=2,
        tickmode="array",
        tickvals=[10 ** (-4 * i) for i in range(5)],
        ticktext=[f"{10**(-4*i):.0e}" for i in range(5)],
        title="logFC absolute difference",
    )
    fig.update_xaxes(row=3, col=1, title="inmoose")
    fig.update_yaxes(row=3, col=1, title="DESeq2", tickangle=-90)
    fig.update_xaxes(row=3, col=2, title="inmoose")
    fig.update_yaxes(row=3, col=2, title="pydeseq2", tickangle=-90)
    fig.update_xaxes(row=3, col=3, title="DESeq2")
    fig.update_yaxes(row=3, col=3, title="pydeseq2", tickangle=-90)

    import plotly.express as px

    fig.data[1].marker.color = px.colors.qualitative.Plotly[2]
    fig.data[2].marker.color = px.colors.qualitative.Plotly[1]

    fig.show()


if __name__ == "__main__":
    run_deseq()
    run_edger()
    run_limma()
    plot_barplots()
```
