## Supplementary material for "Differential Expression Analysis with InMoose, the Integrated Multi-Omic Open-Source Environment in Python": run_DESeq2.R

```
library("DESeq2")

# Get all args from the command line
args = commandArgs(trailingOnly=TRUE)

dataset = args[1]
factors = strsplit(args[2], split = ",")[[1]]
contrast = strsplit(args[3], split = ",")[[1]]

# Load files
clinical_df <- read.csv(paste(dataset, "clinical_df.csv", sep="/"), row.names=1)
molecular_df <- read.csv(paste(dataset, "gene_expression_df.csv", sep="/"), row.names=1)

# Reorder and make sure the order is good
samples_intersection <- intersect(colnames(molecular_df), row.names(clinical_df))
molecular_df <- molecular_df[, samples_intersection]

clinical_columns <- colnames(clinical_df)
clinical_df <- as.data.frame(clinical_df[samples_intersection,],
                             row.names=samples_intersection)
colnames(clinical_df) <- clinical_columns

# Create design matrix
formula <- "~ 1"
formula <- paste0(formula, paste0(" + ", factors, collapse=""))

# Create DESeq2 object
dds <- DESeqDataSetFromMatrix(countData=molecular_df,
                              colData=clinical_df,
                              design=as.formula(formula))

# Remove very lowly expressed genes
##keep = rowSums(counts(dds) >= 10) >= 2
##dds <- dds[keep,]

# Launching diffexp
dds <- DESeq(dds)

# Formatting results
res <- results(dds, contrast=contrast)

write.csv(res, paste(dataset, "deseq2.csv", sep="_"))
```
