## Supplementary material for "Differential Expression Analysis with InMoose, the Integrated Multi-Omic Open-Source Environment in Python": run_limma.R

```
library("limma")

# Get all args from the command line
args = commandArgs(trailingOnly=TRUE)

dataset = args[1]
factors = strsplit(args[2], split = ",")[[1]]
contrast = strsplit(args[3], split = ",")[[1]]

# Load files
clinical_df <- read.csv(paste(dataset, "clinical_df.csv", sep="/"), row.names=1)
molecular_df <- read.csv(paste(dataset, "gene_expression_df.csv", sep="/"), row.names=1)

# Reorder and make sure the order is good
samples_intersection <- intersect(colnames(molecular_df), row.names(clinical_df))
molecular_df <- molecular_df[, samples_intersection]

clinical_columns <- colnames(clinical_df)
clinical_df <- as.data.frame(clinical_df[samples_intersection,],
                             row.names=samples_intersection)
colnames(clinical_df) <- clinical_columns

# Create design matrix
formula <- "~ 0"
formula <- paste0(formula, paste0(" + ", factors, collapse=""))
design <- model.matrix(as.formula(formula), data=clinical_df)

for (col in colnames(design)) {
  colnames(design)[colnames(design) == col] <- gsub(" ", "_", gsub("-", "_", gsub(",", "_", col)))
}
fit1 <- lmFit(molecular_df, design)
tt <- paste(paste0(contrast[[1]], gsub(" ", "_", contrast[[3]])),
            paste0(contrast[[1]], gsub(" ", "_", contrast[[2]])), sep="-")
contrast_matrix <- makeContrasts(tt, levels=design)

# Fit with contrast
data_fit_con <- contrasts.fit(fit1, contrast_matrix)
fit1_ebay <- eBayes(data_fit_con)
top_table <- topTable(fit1_ebay, n = Inf, coef = tt)

write.csv(top_table, paste(dataset, "limma.csv", sep="_"))
```
